# PhyloAug: An Evolutionary and Structure-Aware Data Augmentation Tool ^*^

**DOI:** 10.1101/2025.09.05.674571

**Authors:** Jack Cole, Heng Yang, Krasimira Tsaneva-Atanasova, Ke Li

**Affiliations:** Department of Computer Science, University of Exeter, EX4 4QF, Exeter, UK; Department of Mathematics and Statistics, University of Exeter, Stocker Rd, EX4 4QJ, Exeter, United Kingdom

## Abstract

Genomic Language Models (GLMs) suffer from the inherent problem of data scarcity, due to the cost, time and complexity of wet-lab experiments. Data augmentation offers a solution; however traditional methods may unintentionally affect the underlying structure or function. By combining evolutionary signals with the RNA secondary structure, augmentations can retain original function, and remain structurally coherent. To implement this, we developed PhyloAug, a structure-aware, evolution-inspired augmentation method grounded in neutral theory. We employ Genomic Foundation Models (GFMs) to accurately perturb RNA sequences, and utilise phylogenetic analysis via PAML to provide site-wise restrictions based on evolutionary principles. These principles are obtained through the identification of evolutionarily neutral sites (sequence positions where mutations are unlikely to alter function), which are concatenated with the predicted (or known) secondary structure. We thereby ensure adherence to the underlying structure whilst enabling biologically plausible variation. To validate the biological validity of our augmentations, we compare our predicted neutral sites with Rfam-annotated conserved regions and assess sequence similarity to the underlying multiple sequence alignments. We next fine-tune GLMs on augmented data, yielding significant performance improvements up to 12.9% MCC, and 17.2% F1-Score.

## 1 Introduction

Genomic Foundation Models (GFMs) are large-scale machine learning models composed of millions to billions of parameters, pre-trained across an extensive corpus of genomic data. GFMs have been highly appraised for their adaptability to similar genomic tasks,, with fine-tuning acting as the transfer layer to allow a general model, pre-trained through unsupervised learning, to be tuned for a specific task on a much smaller set of supervised data. A wave of innovation has recently demonstrated GFMs potential to decipher the language of genomics, DNA, RNA and proteins, with huge successes in protein structure modelling with AlphaFold [9], Evo 2 for DNA language modelling [3], and the identification of new translation-associated motifs in plant RNA with PlantRNA-FM [35]. However, despite the incredible achievements, the effectiveness of GFMs remains fundamentally limited by the scarcity of high-quality genomic data, particularly when applied to downstream tasks.

Benchmark datasets aim to provide highly accurate representations for genomic data, as to ensure results are representative of real-world data. Prominent genomic benchmarks such as OmniGenBench [32] and BEACON [26] provide curated, diverse genomic datasets. However, existing benchmarks must rely on biological laboratories to verify data, where rigorous pre-processing and data validation techniques are applied with diverse sequencing technologies and wet-lab experimentation. It is difficult to obtain verified labels, with biologically complex tasks, such as structural annotation [31] and functional annotation of long non-coding RNA [6, 19] requiring significant fees and extensive biological expertise to ensure accurate results. It is for this purpose that GFMs are highly sought after to predict the outcome, as to minimise the time and costs associated, however without the original data to fine-tune the GFM, we are unable to obtain accurate results.

This cycle can be broken by the usage of data augmentation, methods which can generate new unseen data using existing datasets, to allow current models to generalise on sparse datasets, and improve performance by oversampling under-represented samples. Although data augmentation is a widely recognised field and has been applied in numerous areas across computer science [17, 23, 29], genomics data is context-dependent [15], and thus widely applied techniques usually applied in natural language processing (NLP), such as random substitution and input reversal, cannot easily be applied to genomics data [27]. Furthermore, unlike common tasks such as sentence-modelling or image-based analysis, we as humans cannot accurately determine the label of genomics data merely by the predictive input, and must rely on biological wet-lab verification. Thus, we must preserve the original data labels during augmentation. Substitution of the real label with a synthetic label may violate a crucial assumption of the augmented dataset, that the augmented data is supported by the true data distribution [28]. Should the augmented data fall outside of the true distribution, it may influence the model to learn or disregard best policies in favour of addressing the incorrect augmentations. These key challenges of genomic modelling motivate our work, PhyloAug, a novel data augmentation methodology leveraging evolutionary biology to amplify the predictive power of genomic foundation models for non-coding RNA-specific tasks.

### 1.1 Mutations in Evolutionary Biology

PhyloAug is motivated through the widely renowned neutral theory within evolutionary biology. This theory asserts that the majority of evolutionary changes at the molecular level, within DNA, RNA and proteomic sequences, are a result of random genetic drift of neutral mutations, rather than Darwinian selection [12]. Whilst this is a highly debated topic within molecular biology [8, 11], it is widely accepted that neutral mutations are a fundamental part of molecular biology. Because these mutations often have no observable phenotypic effect, they provide a biologically sound method for inducing variation within training data through data augmentation without disrupting the underlying functional signals.

Neutral mutations in coding regions, especially those that do not change the resulting protein, are well understood and often used as benchmarks in evolutionary studies, such as the McDonald–Kreitman test [5]. However, while neutral changes also occur frequently in non-coding RNAs, it is much harder to differentiate between mutations that affect function and neutral mutations. This is because non-coding RNAs lack a corresponding amino-acid, thereby making it more difficult to detect the effects of mutations. Many non-coding RNAs, particularly long non-coding RNAs, are believed to evolve through nearly neutral processes [20], where most variants appear “noisy” due to their selective impact being too small to clearly distinguish from random drift. In practice, identifying functional sites in ncRNAs often requires using a combination of structural conservation, covariation patterns, and sequence conservation, rather than relying on simple sequence conservation as in coding RNA. In our work, we predict the neutral mutations using structural conservation estimated through Rfam covariance model-based alignments and the RNA secondary structure, and utilise PAML to identify sequence conservation patterns. We motivate PhyloAug as an augmentation methodology that aims to utilise these neutral mutations within augmentation. We integrate this evolutionary data by obtaining neutral site estimates through computationally-derived evolution.

### 1.2 RNA Structure in GFMs

As well as utilising evolutionary principles, we also aim to incorporate structural data within our pipeline, as to preserve RNA-Protein interactions and prevent the model from learning impeding sequences that may obscure the original function. Previous work such as OmniGenome [33] and RNAErnie [30] has demonstrated that incorporation of the RNA secondary structure can provide additional context, such as vital motifs within the RNA that must be preserved. Thus, by incorporating the secondary structure in our pipeline, we can identify key structural motifs important to function.

Many RNAs released contain their secondary structure, however if the secondary structure cannot be obtained, we utilise ViennaRNA [18], a secondary structure prediction method based on thermodynamic principles, to estimate the true secondary structure. This guides our augmentation process to minimise disruption through avoiding computationally-derived mutations for the predicted secondary structure, preserving the original function.

### 1.3 Our Contributions

Our work studies new techniques for genomic data augmentation, with the goal to restrict augmentations to the distribution of the original dataset through evolutionary and structural constraints. We further motivate our work in the Examining Methodology Effectiveness subsection of Experiments, by performing an experimental investigation into how neutral mutations can prevent the adjustment of conserved nucleotides in RNA sequences, utilising the Rfam-annotated families as our ground-truth. This insight motivates our major contribution, the data augmentation pipeline established with evolutionary and structural constraints. We describe the core of our methodology in the Overall Pipeline section of Methodology, and discuss the incorporation of phylogenetic analysis and statistical testing to ensure robust and replicable results. We further outline our neutral site identification pipeline in fig. 1, and our overall data pipeline. In the Model Performance subsection of Experiments, we present and discuss an evaluation of our augmentation strategy when applied to key RNA-based tasks and GFM model architectures, demonstrating consistent improvement across state-of-the-art models.

**Figure 1.**
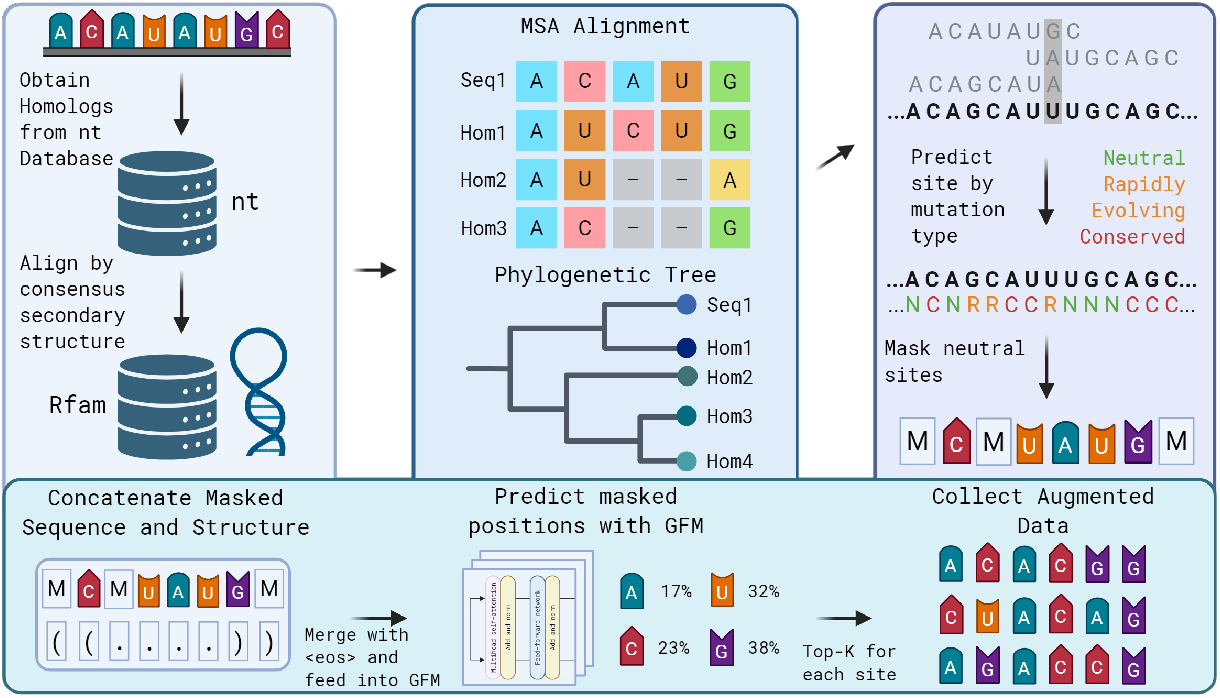
Overview of the augmentation pipeline. The process begins with the retrieval of homologous sequences from the NCBI nt database. Sequence-only homologs, homologs that do not belong to the same structural family, are filtered out using annotations from the Rfam database. A multiple sequence alignment is then constructed using MAFFT, and a phylogenetic tree is constructed using FastTree. These are then fed into PAML, where we employ ancestral sequence reconstruction and the estimation of site-specific evolutionary rates. Based on these rates, each site is classified as conserved, rapidly evolving, or putatively neutral. Only sites inferred to be neutral are masked. The masked sequence is then concatenated with its corresponding secondary structure and passed to the genomic foundation model, which perturbs the masked regions to generate the final augmented sequence set.

## 2 Related Work

The primary challenge of limited high-quality genomic data has led to several approaches aimed at either generating new data through data augmentation, or enriching existing data with biological context. These methods broadly fall into two categories: data augmentation and the direct integration of evolutionary information into model architectures. Previous data augmentation methodologies include [14], who utilise Generative Adversarial Models (GANs) to learn the original distribution of the data, and perturb the data to create augmentations. Despite success on their experimental datasets, methodological analysis revealed a low recall scores, thereby suggesting that the models were unable to learn the true underlying data distribution. An alternative work is EvoAug [16], which utilises evolution-inspired techniques (deletion, mutation, translocation, etc.) to create an augmented training set for deep learning models to be trained on, after which they are fine-tuned on the original training dataset. However, this method cannot be applied to GFMs, where it may take months to complete pre-training, and requires deep biological knowledge to ensure the underlying data structure (RNA secondary structure and conserved nucleotides) is preserved during augmentation.

### 2.1 Leveraging Evolutionary Information in GFMs

Previous work primarily focuses on the usage of Multiple Sequence Alignment (MSA) or Phylogenetic information through Phylogenetic trees. MSA is used to align DNA or RNA sequences that are evolutionary similar, as to uncover the evolutionary mutations that have occurred per sequence. Phylogenetic trees are branching diagrams that illustrate the evolutionary relationship of a single or group of organisms. MSA is often termed as a “horizontal approach”, and Phylogenetic Analysis as a “vertical approach”, in which the evolutionary structure of the sequences is preserved [21], although just incorporating the phylogenetic tree is not enough to reach this vertical approach, and we must further utilise ancestral reconstruction tools such as PAML (Phylogenetic Analysis by Maximum Likelihood) [34]. Ancestral reconstruction re-creates the original sequences before evolutionary mutation occurred, thereby allowing us to make direct comparisons and accurately determine the types of mutations that occurred at each site of the nucleotide sequence. Whilst PAML is traditionally used for coding RNA, previous studies have demonstrated the use of the BaseML function for phylogenetic analysis of non-coding RNA [7]. This provides a clear and accurate framework for the identification of neutral mutations by utilising the evolutionary context.

GFMs generally focus on the incorporation of MSA and phylogenetic information within the model pipeline, as a way to directly inject evolutionary information directly into the model. There have been several approaches, such as RNA-MSM [36], a GFM pre-trained on RNA-MSA data, and the MSATransformer [25] proposed for Protein Language Models, enabling pre-training across a huge variety of MSA data. However, whilst incorporating MSA data has proved to be beneficial, GPN-MSA [2] suggests the incorporation of evolutionary information alongside the MSA can further improve performance, as through aligning the MSA with the same gene across 100 vertebrate species, they achieve state-of-the-art performance in variant effect prediction. [1] and [37] are further recent examples of integrating phylogenetic information within genomic models. CSFold establishes several key limitations of MSA, such as the reliance on the most common nucleotide, rather than establishing a clear evolutionary trend and pattern, and further utilises PAML and statistical tests to provide insight into the evolutionary trends of the data. PhyloGPN establishes a novel training paradigm where the training loss is used to model the evolution of aligned nucleotides given a phylogenetic tree, thus training the model to inherently understand nucleotide evolution.

### 2.2 Overview

These previous works have established that adding additional evolutionary-based information through MSA or Phylogenetic Trees may improve algorithm performance, however the method including this information varies greatly, and is algorithm-specific. We propose a one-size-fits-all solution for noncoding RNA, which can be applied to any gFM through fine-tuning, utilising the evolutionary and structural information through data augmentation.

## 3 Methodology

### 3.1 Overall Pipeline

To ensure that we do not encounter the problem of damaging the sequence function or structure discussed previously, we introduce a neutral masking methodology pipeline using established bioinformatics tools. We use grounded biological theory to ensure that the nucleotides we mask minimally impact biological function or structure, and therefore result in an augmented set of sequences that are closely tied to the true dataset, and still fit the original labels. Our overall approach is as follows; gather homologous sequences using BLASTN and the nt database, utilise biological pipelines to establish neutral positions, mask them, recover masking percentage to a set threshold, and feed in the masked sequences to the GFM. However, the tools used to identify neutral positions do not necessarily consider the secondary structure of the RNA. In order to adhere to this, we incorporate two additional features. Firstly, we adjust our data to ensure all data incorporated for neutral position identification uses Rfam-family aligned data, thus aligning all of our sequences through both sequence and structure. We further concatenate the secondary structure into our GFM, to allow the model to consider the pseudoknot-free pairings that may affect our results. Lastly, we prevent motifs that cannot be considered by our pipeline, but are still present in our secondary structure annotation (e.g., pseudoknots) from being masked, as to preserve the RNA structure as closely as possible. With this method, we have created augmentations that adhere to the evolutionary and structural features of our underlying data.

#### 3.1.1 Identifying Neutral Mutations

A neutral mutation in is a change in a genomic sequence that has no effect on function or organism fitness. In coding RNA, synonymous substitutions (base changes that do not alter the amino acid) are assumed to be neutral mutations, as leaving the protein sequence unaffected will result in the function being unaffected also [4]. Although non-synonymous substitutions alter the amino acid sequence, they can still be neutral if they do not impact the protein’s structure or activity. In practice, neutral evolution is estimated through the ratio of non-synonymous and synonymous substitutions rates, defined as *dN/dS*, and denoted as *ω*. When *ω* is 1, non-synonymous changes occur at the same rate as synonymous changes, which is the rate at which neutral evolution happens in evolution. This distinction is crucial, as, rather than following previous work which simply works to change nucleotides without changing the protein codons, we utilise *ω* to further predict non-synonymous substitutions that are neutral. To do this, we utilise PAML, which reconstructs the ancestral sequences using the MSA and phylogenetic tree, and estimates evolutionary parameters like *ω* during phylogenetic analysis.

#### 3.1.2 Predicting Rapidly Evolving Sites

Whilst identifying neutral mutations is a well known and explored field in the realm of coding RNA, for non-coding RNA (ncRNA), this problem is not so trivial. In coding RNA, a mutation can directly affect the amino-acid codons, thereby it is much easier to detect a direct change. Unlike coding RNA, ncRNAs do not have codons, and function through structural and regulatory roles, meaning mutations cannot be directly assessed through the amino-acid chain. Furthermore, mutations that do not alter the sequence function may still impact secondary structure, RNA-protein interactions, or expression. Therefore identifying neutral variation in ncRNA requires additional information, including the folded secondary RNA structure, and the usage of comparative genomics to identify nucleotides that are susceptible to change.

We begin by individually selecting each sequence from the training dataset, and performing an exhaustive homology search using NCBI’s nt database, which contains over 116M RNA or DNA sequences. This allows us to enrich the dataset by integrating evolutionary information from closely related sequences, thereby forming the basis of our Multiple Sequence Alignment (MSA). To ensure the sequences are homologically related, we utilise an e-value of 1*e −* 5, ensuring that homologous sequences that have undergone significant change can be utilised. After collecting our homologs, and subsequently removing any duplicates, we utilise Rfam to align our sequences by structural family. By utilising this method, we ensure our homologs are both sequentially and structurally aligned with our underlying sequence from the training dataset. Next, MAFFT [10] is used to computationally align the sequences through MSA, by accounting for evolutionary events such as insertions, deletions, and substitutions that have accumulated over extensive evolutionary timescales. By using homologs that share a common evolutionary ancestry, we can enrich our models with evolutionary information, such as the conserved nucleotides that enhance or diminish function within the sequence, and identification of neutral mutations that do not affect function. However, phylogenetic analysis of just the MSA has key limitations, such as obscuring compensatory substitutions due to prioritising column-wise frequency over evolutionary information.

Thus, in parallel with MSA, we construct a phylogenetic tree based on the aligned RNA sequences within each homologous family using FASTTREE [24]. Our phylogenetic analysis tool, PAML, is known to be inaccurate when constructing phylogenetic trees, thus we create our own using FASTTREE. The incorporation of a phylogenetic tree provides a framework for understanding the evolutionary trajectory of RNA families, revealing ancestral lineages and pinpointing evolutionary events obscured by MSA. Next, we combine the aligned sequences and phylogenetic tree with PAML (Phylogenetic Analysis by Maximum Likelihood), to perform ancestral sequence reconstruction. This step utilises maximum likelihood methods to infer probable ancestral RNA sequences at internal nodes of the phylogenetic tree.

Rather than directly utilising reconstructed ancestral sequences, we opt to analyse the evolutionary patterns to rule out conserved and fast-evolving mutations. In particular, we employ PAML’s baseml tool to estimate site-specific substitution rates from the provided MSA and phylogenetic tree. Sites that exhibit very low substitution rates are inferred to be conserved, likely due to structural or regulatory importance, while rapidly evolving sites may indicate natural selection or adaptation. Both types are assumed to be functionally important and are therefore excluded from our candidate set of neutral mutations, as is in-line with previous phylogenetic analysis of non-coding RNA [13, 22]. To identify these constrained positions, we fit nucleotide substitution models with rate variation across sites using a discrete gamma distribution and empirical Bayes approaches. From this, we obtain relative rate estimates for each site. These form the basis of the constrained masking strategy, where we exclude conserved or rapidly evolving sites, retaining only sites with moderate substitution rates as likely neutral. This filtering approach allows us to avoid perturbing functionally important regions during downstream tasks such as data augmentation or model training.

We aim to mask roughly 15% of the sequence positions as neutral, although as each sequence is different, some variability must be allowed. There are two key challenges, over and under-identification of sites. Should we over-identify sites, we simply utilise stricter filtering, or unmask nucleotides at the end points of our neutral estimates until we reach our threshold of masked nucleotides (15%). Alternatively, under-identifying sites may not provide enough sequence diversity for our augmentation model, thus we utilise a cut-off point. If at least 10% of nucleotides are not masked, the sequence is excluded from augmentation. As we have constrained our site-wise selection for masking, we thereby reduce the impact that our augmentations will have on the underlying biologically processes and functions of the ncRNA sequence.

### 3.2 Combining Neutral Positions with gFMs

A key motivation behind the restriction to a mere 15% threshold of masked nucleotides in the masking fully strategy is the understanding that GFMs require enough sequential context. Furthermore, our masking strategy does not fully consider the RNA secondary structure when identifying rapidly evolving nucleotides, although we do consider the structural family in order to accurately reconstruct the sequence. Thereby, just providing the masked sequence alone could result in accidental changes within the secondary structure. To solve these limitations, we concatenate the secondary structure label with the RNA sequence, to inject secondary context into the model, thereby minimising substitutions that may modify the secondary structure. We selected OmniGenome as our GFM for perturbing the RNA sequences, as OmniGenome was pre-trained with multiple objectives, Masked-Language-Modeling with a mask percentage of 15%, and with concatenated RNA secondary structures and sequences. As this model has been pre-trained with 54.2B tokens using these direct objectives, it is therefore the best fit to ensure best performance.

To accurately fill in the masked positions, we utilise a top-k approach, where only the top nucleotides will be considered, dependent on the softmax output of the model. A similar distribution of likelihoods across all nucleotides suggests all nucleotides are compatible for a mutation, however in the case of base-pairs being compatible, the mutation may be much more restrictive. Once the augmentations are complete for the training set, we merge the augmented sequences with the original dataset, and fine-tune our models.

### 3.3 MSA-Only based Approach

Notably in our pipeline, we split the type of mutation into three separate types, neutral, rapidly evolving and conserved, where according to biological theory, neutral evolutions are sites that frequently change, however, if the site is rapidly evolving, it may signify an importance to enhancing or decaying function. Furthermore, if a site rarely changes, it may signify an important site that is conserved. Through this description, it is possible to estimate these categories using the Multiple Sequence Alignment file alone, although doing so is known to be unreliable. To prove the effectiveness of phylogenetic analysis, we incorporate this as a comparator, as to provide further information on the importance of each part of the pipeline.

## 4 Experiments

### 4.1 Examining Methodology Effectiveness

Whilst we know based on biological theory that for coding RNA, our pipeline will effectively identify neutral candidates, for non-coding RNA, there is no established “best way” to locate neutral RNA. Thus, we investigate the effectiveness of this critical part of our method as to ensure the reliability and effectiveness of our method on ncRNA. This experiment aims to identify the effectiveness of our method to avoid conserved nucleotides, key sites where adjusting them could influence functional motifs. To accomplish this, we first select 6 diverse Rfam families, RF00001, RF00005, RF00051, RF00163, RF00906 and RF03160. RF03160, RF00051, RF00001 and RF00005 represent Rfam families with a large amount of sequences, with 50 homologs used to build the conserved nucleotide space. RF00163 and RF00906 contain 20 homologs, as to represent families with a less known set of conserved nucleotides. We utilise a maximum of 50 sequences within the Rfam-established MSA built for the families, and put our randomly selected set of sequences through our pipeline. We then measure the success of our method by calculating overlap between the conserved sites and our masked sites, where the conserved positions are obtained from the original Rfam-annotated data. To further establish the effectiveness of the incorporation of the Rfam-family alignment, we remove the Rfam part of our pipeline and show only the results of the phylogenetic analysis section of our method.

### 4.2 Results

We find that PhyloAug with all parts of our method, phylogenetic analysis and rfam-family alignment, performs the best, with a clash rate of merely 0.073% with conserved nucleotides. We further find that all methods, even including our naive MSA method, outperform random selection. We find that each part of our methodology linearly increases the effectiveness, although the incorporation of the Rfam-family alignment provides the most significant increase in effectiveness. These results demonstrate the ability of our method to improve performance a variety of downstream tasks, by avoiding sites vital to both function and structure.

**Table 1:**
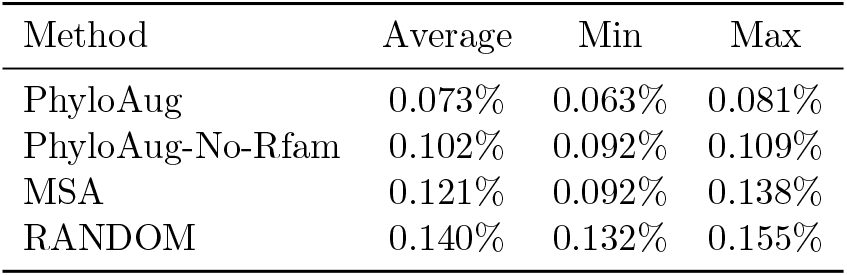
Average number of conserved nucleotides masked with each augmentation method. PhyloAug represents the full pipeline described in Overall Pipeline, PhyloAug-No-Rfam represents the pipeline without using Rfam to predict the conserved nucleotides, MSA represents the MSA-only methodology previously described, and Random represents masking based on purely random nucleotides with no constraints. A masking rate of 15% was used for each method.

### 4.3 Improving Structural Prediction

#### 4.3.1 Experimental Design

To ensure the validity of our approach, we run several experiments on publicly available datasets. We split our experiments into Non-Coding and Coding RNA, as per the methodology for obtaining neutral mutations described previously. We utilise the three standard structural prediction datasets, ArchiveII, bpRNA, and rnastralign, as consistent with previous analysis and benchmarking methods. To evaluate the performance, we utilise the standard F1-Score, and combine it with Matthew’s Correlation Coefficient (MCC), as to further evaluate the robustness of our model performance. F1-Score does not evaluate true negatives, thus by including MCC, we also evaluate the negative prediction aspect of our models.

#### 4.3.2 Results

When testing our augmentation method for non-coding RNA, we find that for all structural prediction tasks, our augmentation methodology improves performance consistently across all models. For Archive2, we find that small models, such as RNA-BERT, which were previously unable to generalise to the sparse dataset has a major increase in both MCC and F1-score. Models with stronger performance, such as RNA-FM and RNA-MSM see a small increase F1-Score, but a comparatively larger increase in MCC, suggesting a reduction in false positives and negatives across model performance. Therefore, augmentations increase the robustness of the models, as well as their predictive accuracy. We see a similar trend in the bpRNA dataset, whereas rnastralign also shows signs of this trend, for top performing models such as RNA-FM and OmniGenome, there is little change in model performance. This is not unexpected however, as the model performance is almost perfect, suggesting that this may be the peak of the model understanding for this dataset.

#### 4.4 Sequence pattern preservation and distribution

In this section we aim to answer how effective our augmentation methodology is at preserving evolutionarily conserved contexts encoded within biological sequences. To achieve this, we utilise the non-coding RNA structural prediction datasets, and assess how closely the augmentations can reproduce the underlying characteristics of the set of homologous sequences. We perform this investigation on each individual RNA within all three ncRNA structural prediction datasets, as to provide a complete and comprehensive evaluation.

#### 4.4.1 Experimental Design

Traditional analysis of RNA motif preservation utilises a simple visual inspection of sequence logos, however due to the large amount of augmented sequences, we cannot show an accurate visual diagram across all data. To preserve the motif-specific interactions across each original RNA and it’s augmented set, we compare the MSA-aligned homologs obtained for each RNA with the augmented set of RNA sequences. To fully understand the overall similarity across the entire training set, we average the Jensen-Shannon Distance (JSD) and Cosine Similarity scores across all augmented sequences as opposed with the MSA-aligned homologs. This gives us an average for how closely the augmented sequence is able to represent the underlying nucleotide distribution across the underlying data proportions. JSD is used to measure the similarity of the nucleotide distributions, and Cosine Similarity measures the overall pattern/shape of the nucleotide frequencies.

**Table 2:**
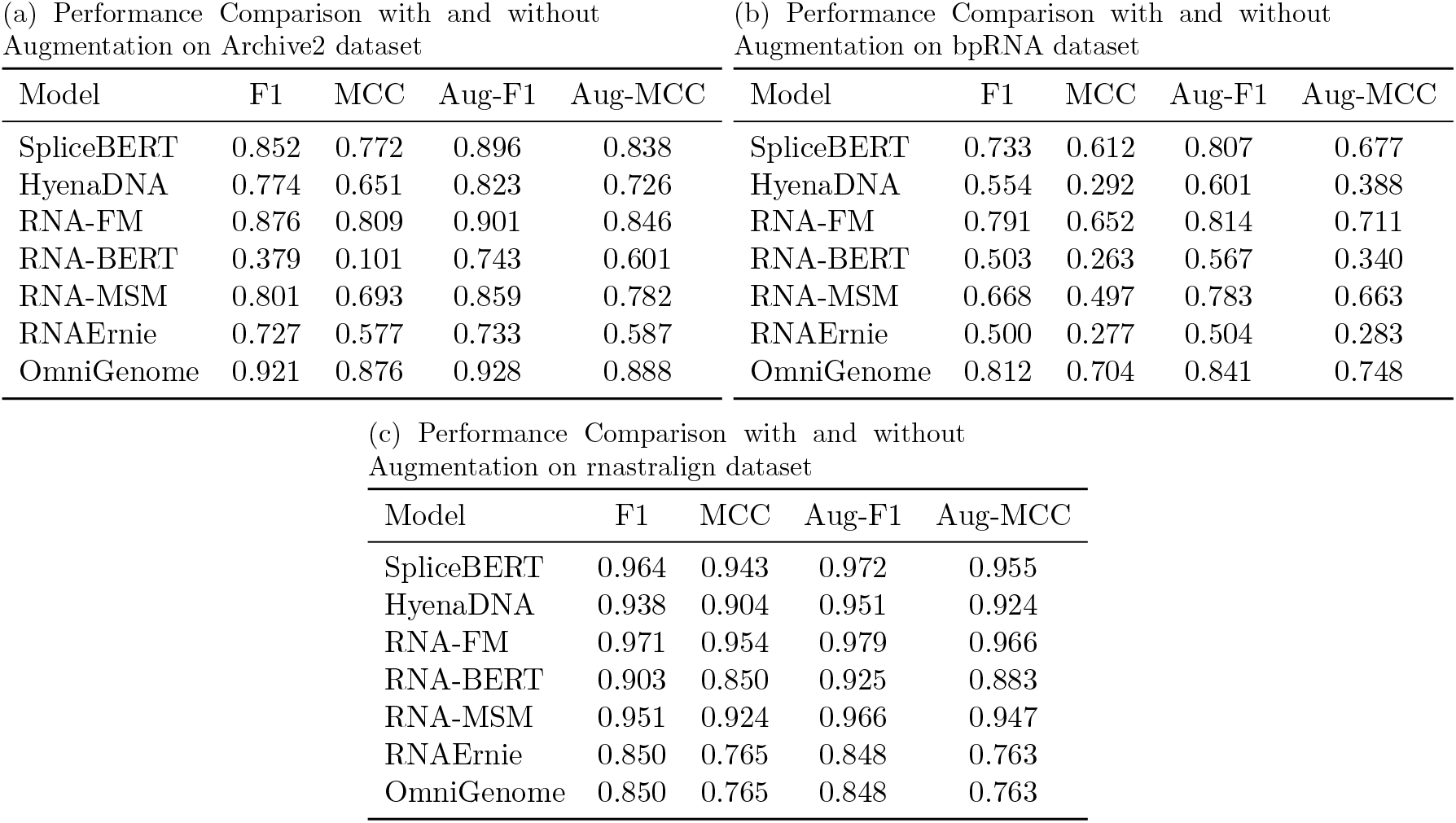
Performance comparison of various models with and without augmentation on different datasets. (a) Performance Comparison with and without Augmentation on Archive2 dataset (b) Performance Comparison with and without Augmentation on bpRNA dataset (c) Performance Comparison with and without Augmentation on rnastralign dataset

| Model | F1 | MCC | Aug-F1 | Aug-MCC |
| --- | --- | --- | --- | --- |
| SpliceBERT | 0.852 | 0.772 | 0.896 | 0.838 |
| HyenaDNA | 0.774 | 0.651 | 0.823 | 0.726 |
| RNA-FM | 0.876 | 0.809 | 0.901 | 0.846 |
| RNA-BERT | 0.379 | 0.101 | 0.743 | 0.601 |
| RNA-MSM | 0.801 | 0.693 | 0.859 | 0.782 |
| RNAErnie | 0.727 | 0.577 | 0.733 | 0.587 |
| OmniGenome | 0.921 | 0.876 | 0.928 | 0.888 |

(b) Performance Comparison with and without Augmentation on bpRNA dataset
| Model | F1 | MCC | Aug-F1 | Aug-MCC |
| --- | --- | --- | --- | --- |
| SpliceBERT | 0.733 | 0.612 | 0.807 | 0.677 |
| HyenaDNA | 0.554 | 0.292 | 0.601 | 0.388 |
| RNA-FM | 0.791 | 0.652 | 0.814 | 0.711 |
| RNA-BERT | 0.503 | 0.263 | 0.567 | 0.340 |
| RNA-MSM | 0.668 | 0.497 | 0.783 | 0.663 |
| RNAErnie | 0.500 | 0.277 | 0.504 | 0.283 |
| OmniGenome | 0.812 | 0.704 | 0.841 | 0.748 |

Table 2: Performance comparison of various models with and without augmentation on different datasets.
| Model | F1 | MCC | Aug-F1 | Aug-MCC |
| --- | --- | --- | --- | --- |
| SpliceBERT | 0.964 | 0.943 | 0.972 | 0.955 |
| HyenaDNA | 0.938 | 0.904 | 0.951 | 0.924 |
| RNA-FM | 0.971 | 0.954 | 0.979 | 0.966 |
| RNA-BERT | 0.903 | 0.850 | 0.925 | 0.883 |
| RNA-MSM | 0.951 | 0.924 | 0.966 | 0.947 |
| RNAErnie | 0.850 | 0.765 | 0.848 | 0.763 |
| OmniGenome | 0.850 | 0.765 | 0.848 | 0.763 |

#### 4.4.2 Results

Our results across all three datasets show a significant improvement for PhyloAug as opposed to MSA-only and random masking. We find that for each level of augmentation, we gradually approach the ground truth, and with only one level of augmentation, the overall result is the largest distance away from the ground truth. This is intuitive as our augmented sequences should represent similar homologs our MSA-aligned data. This further demonstrates the usefulness of a large set of augmentations, as with increasing augmentations, we draw closer to the ground truth. We find that the full PhyloAug pipeline results in the closest evolutionary distance from the ground truth, with random masking being the furthest. There is a significant difference between PhyloAug and the alternative methods, of which the gap is maintained as the augmentation level rises. This thereby proves our empirical result, being that the more augmentations, the better overall performance, with reducing returns. The low JSD values demonstrate that the nucleotide distributions of our augmented sequences are closely aligned with the original MSA homologs, and our high cosine similarity shows a similar overall pattern/shape of nucleotides. We thereby demonstrate empirically through performance and evolutionary distance the effectiveness of our method.

**Table 3:** JSD and Cosine Similarity results for three datasets.

|  | Dataset / Augs | Phylo Masking |  | MSA-only Masking |  | Random Masking |  |
| --- | --- | --- | --- | --- | --- | --- | --- |
|  |  | JSD | Cosine | JSD | Cosine | JSD | Cosine |
| **Archive2** | 1 | 0.2431 | 0.7654 | 0.2740 | 0.7349 | 0.2896 | 0.7205 |
|  | 2 | 0.2422 | 0.7674 | 0.2732 | 0.7361 | 0.2810 | 0.7291 |
|  | 4 | 0.2414 | 0.7690 | 0.2728 | 0.7369 | 0.2745 | 0.7350 |
|  | 8 | 0.2410 | 0.7694 | 0.2726 | 0.7373 | 0.2711 | 0.7379 |
|  | 12 | 0.2402 | 0.7702 | 0.2726 | 0.7375 | 0.2703 | 0.7386 |
| **bpRNA** | 1 | 0.2279 | 0.7799 | 0.2558 | 0.7499 | 0.2883 | 0.7202 |
|  | 2 | 0.2264 | 0.7821 | 0.2550 | 0.7533 | 0.2780 | 0.7303 |
|  | 4 | 0.2242 | 0.7864 | 0.2544 | 0.7448 | 0.2708 | 0.7360 |
|  | 8 | 0.2236 | 0.7879 | 0.2540 | 0.7456 | 0.2667 | 0.7393 |
|  | 12 | 0.2228 | 0.7887 | 0.2536 | 0.7466 | 0.2653 | 0.7404 |
| **rnastralign** | 1 | 0.2363 | 0.7838 | 0.2582 | 0.7619 | 0.2665 | 0.7577 |
|  | 2 | 0.2362 | 0.7847 | 0.2564 | 0.7658 | 0.2613 | 0.7634 |
|  | 4 | 0.2358 | 0.7855 | 0.2548 | 0.7677 | 0.2576 | 0.7666 |
|  | 8 | 0.2353 | 0.7861 | 0.2540 | 0.7689 | 0.2555 | 0.7685 |
|  | 12 | 0.2346 | 0.7863 | 0.2536 | 0.7694 | 0.2549 | 0.7689 |

## 5 Conclusions

In this work, we introduced PhyloAug, a data augmentation method incorporating both evolutionary and structural information to improve Genomic Foundation Models on downstream tasks. We first empirically demonstrate the effectiveness of incorporating evolutionary information into the data augmentation process, and establish the importance of the key sections of our pipeline. We further provide comparisons to naive methods such as MSA-only, and random-based methods. Comprehensive experiments illustrate the effectiveness of PhyloAug within non-coding RNA, and empirically demonstrates improved downstream performance across several key ncRNA-based tasks. We hope that PhyloAug will be utilised as a tool to enhance the performance and robustness of current GFMs, and allow GFMs to generalise on small, task specific datasets prevalent in genomic downstream tasks.

## Acknowledgment

This work was supported in part by the UKRI Future Leaders Fellowship under Grant MR/S017062/1 and MR/X011135/1; in part by NSFC under Grant 62376056 and 62076056; in part by the Royal Society under Grant IES/R2/212077; in part by the EPSRC under Grant 2404317; in part by the Kan Tong Po Fellowship (KTP*\*R1*\*231017); and in part by the Amazon Research Award and Alan Turing Fellowship.

## References

[1] Carlos Albors, Jianan Canal Li, Gonzalo Benegas, Chengzhong Ye, and Yun S. Song. A phylogenetic approach to genomic language modeling. In Research in Computational Molecular Biology: 29th International Conference, RECOMB 2025, Seoul, South Korea, April 26–29, 2025, Proceedings, 2025.

[2] Gabriel Benegas, Clara Albors, Alvin J. Aw, Chuan Ye, and Y. S. Song. A dna language model based on multispecies alignment predicts the effects of genome-wide variants. Nature Biotechnology, 2025.

[3] Garyk Brixi, Matthew G Durrant, Jerome Ku, Michael Poli, Greg Brockman, Daniel Chang, Gabriel A Gonzalez, Samuel H King, David B Li, Aditi T Merchant, Mohsen Naghipourfar, Eric Nguyen, Chiara Ricci-Tam, David W Romero, Gwanggyu Sun, Ali Taghibakshi, Anton Vorontsov, Brandon Yang, Myra Deng, Liv Gorton, Nam Nguyen, Nicholas K Wang, Etowah Adams, Stephen A Baccus, Steven Dillmann, Stefano Ermon, Daniel Guo, Rajesh Ilango, Ken Janik, Amy X Lu, Reshma Mehta, Mohammad R.K. Mofrad, Madelena Y Ng, Jaspreet Pannu, Christopher Re, Jonathan C Schmok, John St. John, Jeremy Sullivan, Kevin Zhu, Greg Zynda, Daniel Balsam, Patrick Collison, Anthony B. Costa, Tina Hernandez-Boussard, Eric Ho, Ming-Yu Liu, Tom McGrath, Kimberly Powell, Dave P. Burke, Hani Goodarzi, Patrick D Hsu, and Brian Hie. Genome modeling and design across all domains of life with evo 2. bioRxiv, 2025.

[4] L. Calderoni, O. Rota-Stabelli, E. Frigato, A. Panziera, S. Kirchner, N. S. Foulkes, and C. Bertolucci. Relaxed selective constraints drove functional modifications in peripheral photoreception of the cavefish P. andruzzii and provide insight into the time of cave colonization. Heredity, 2016.

[5] Jane Charlesworth and Adam Eyre-Walker. The mcdonald–kreitman test and slightly deleterious mutations. Molecular Biology and Evolution, 2008.

[6] Abhay Chowdhary, Venkata Satagopam, and Reinhard Schneider. Long non-coding rnas: Mechanisms, experimental, and computational approaches in identification, characterization, and their biomarker potential in cancer. Frontiers in Genetics, 2021.

[7] Zhirui Hu, Timothy B Sackton, Scott V Edwards, and Jun S Liu. Bayesian detection of convergent rate changes of conserved noncoding elements on phylogenetic trees. Molecular Biology and Evolution, 2019.

[8] Jeffrey D. Jensen, Bret A. Payseur, Wolfgang Stephan, Charles F. Aquadro, Michael Lynch, Deborah Charlesworth, and Brian Charlesworth. The importance of the neutral theory in 1968 and 50 years on: A response to kern and hahn 2018. Evolution, 2019.

[9] John Jumper, Richard Evans, Alexander Pritzel, Tim Green, Michael Figurnov, Olaf Ronneberger, Kathryn Tunyasuvunakool, Russ Bates, Augustin Žídek, Anna Potapenko, et al. Highly accurate protein structure prediction with alphafold. Nature, 2021.

[10] Kazutaka Katoh and Daron M. Standley. Mafft multiple sequence alignment software version 7: Improvements in performance and usability. Molecular Biology and Evolution, 2013.

[11] Andrew D. Kern and Matthew W. Hahn. The neutral theory in light of natural selection. Molecular Biology and Evolution, 2018.

[12] Motoo Kimura. Evolutionary rate at the molecular level. Nature, 1968.

[13] Jennifer L. Knies, Kristen K. Dang, Todd J. Vision, Noah G. Hoffman, Ronald Swanstrom, and Christina L. Burch. Compensatory evolution in rna secondary structures increases substitution rate variation among sites. Molecular Biology and Evolution, 2008.

[14] Alice Lacan, Michele Sebag, and Blaise Hanczar. Gan-based data augmentation for transcriptomics: survey and comparative assessment. Bioinformatics, 2023.

[15] H. Lee, U. Ozbulak, H. Park, K. Lee, and H. Yang. Assessing the reliability of point mutation as data augmentation for deep learning with genomic data. BMC Bioinformatics, 2024.

[16] Nicholas Keone Lee, Ziqi Tang, Shushan Toneyan, and Peter K. Koo. Evoaug: improving generalization and interpretability of genomic deep neural networks with evolution-inspired data augmentations. Genome Biology, 2023.

[17] Bohan Li, Yutai Hou, and Wanxiang Che. Data augmentation approaches in natural language processing: A survey. AI Open, 2022.

[18] Ronny Lorenz, Stephan H. Bernhart, Christian Höner zu Siederdissen, Hakim Tafer, Christoph Flamm, Peter F. Stadler, and Ivo L. Hofacker. Viennarna package 2.0. Algorithms for Molecular Biology, 2011.

[19] John S. Mattick, Pedro P. Amaral, Piero Carninci, et al. Long non-coding rnas: definitions, functions, challenges and recommendations. Nature Reviews Molecular Cell Biology, 2023.

[20] John S. Mattick, Pedro P. Amaral, Piero Carninci, et al. Long non-coding rnas: definitions, functions, challenges and recommendations. Nature Reviews Molecular Cell Biology, 2023.

[21] Rainer Merkl and Reinhard Sterner. Ancestral protein reconstruction: techniques and applications. Biological Chemistry, 2016.

[22] Sonja Meyer and Arndt von Haeseler. Identifying site-specific substitution rates. Molecular Biology and Evolution, 2003.

[23] Alhassan Mumuni and Fuseini Mumuni. Data augmentation: A comprehensive survey of modern approaches. Array, 2022.

[24] Morgan N. Price, Paramvir S. Dehal, and Adam P. Arkin. Fasttree 2 – approximately maximumlikelihood trees for large alignments. PLoS ONE, 2010.

[25] Roshan Rao, Jason Liu, Robert Verkuil, Joshua Meier, John F. Canny, Pieter Abbeel, Tom Sercu, and Alexander Rives. Msa transformer. In Proceedings of the 38th International Conference on Machine Learning, Proceedings of Machine Learning Research, 2021.

[26] Yuchen Ren, Zhiyuan Chen, Lifeng Qiao, Hongtai Jing, Yuchen Cai, Sheng Xu, Peng Ye, Xinzhu Ma, Siqi Sun, Hongliang Yan, Dong Yuan, Wanli Ouyang, and Xihui Liu. Beacon: Benchmark for comprehensive rna tasks and language models, 2024.

[27] M. Sanabria, J. Hirsch, P.M. Joubert, K. Kreeger, and M. Mele. Dna language model grover learns sequence context in the human genome. Nature Machine Intelligence, 2024.

[28] Han Shao, Omar Montasser, and Avrim Blum. A theory of pac learnability under transformation invariances. In Proceedings of the 36th International Conference on Neural Information Processing Systems, 2022.

[29] Brandon Trabucco, Kyle Doherty, Max Gurinas, and Ruslan Salakhutdinov. Effective data augmentation with diffusion models, 2023.

[30] Nanyi Wang, Jiayi Bian, Yichen Li, Lihua Zhang, Qian Liu, Zhihao Zhou, Yu Zhang, Zhihua Wei, Jingyi Wang, Xuehai Ren, and Ting Liu. Multi-purpose rna language modelling with motif-aware pretraining and type-guided fine-tuning. Nature Machine Intelligence, 2024.

[31] Kyle E. Watters, Alexander M. Yu, Eric J. Strobel, Anthony H. Settle, and Julius B. Lucks. Characterizing rna structures in vitro and in vivo with selective 2’-hydroxyl acylation analyzed by primer extension sequencing (shape-seq). Methods, 2016.

[32] Heng Yang, Jack Cole, Yuan Li, Renzhi Chen, Geyong Min, and Ke Li. Omnigenbench: A modular platform for reproducible genomic foundation models benchmarking. 2024.

[33] Heng Yang, Renzhi Chen, and Ke Li. Bridging sequence-structure alignment in rna foundation models. In Proceedings of the AAAI Conference on Artificial Intelligence, 2025.

[34] Ziheng Yang. PAML 4: Phylogenetic analysis by maximum likelihood. Molecular Biology and Evolution, 2007.

[35] Haopeng Yu, Heng Yang, Wenqing Sun, Zongyun Yan, Xiaofei Yang, Huakun Zhang, Yiliang Ding, and Ke Li. An interpretable rna foundation model for exploring functional rna motifs in plants. Nature Machine Intelligence, 2024.

[36] Yikun Zhang, Mei Lang, Jiuhong Jiang, Zhiqiang Gao, Fan Xu, Thomas Litfin, Ke Chen, Jaswinder Singh, Xiansong Huang, Guoli Song, Yonghong Tian, Jian Zhan, Jie Chen, and Yaoqi Zhou. Multiple sequence alignment-based RNA language model and its application to structural inference. Nucleic Acids Research, 2023.

[37] Jiren Zhou, Jiajia Xu, Jiayu Wen, and Brian John Parker. CS-FOLD: Advancing RNA Structure Predictions through Phylogenetic Modelling of Compensatory Mutations in Deep Neural Networks. bioRxiv, 2025.

